# A high-quality reference genome assembly of the saltwater crocodile, *Crocodylus porosus*, reveals patterns of selection in Crocodylidae

**DOI:** 10.1101/767939

**Authors:** Arnab Ghosh, Matthew G. Johnson, Austin B. Osmanski, Swarnali Louha, Natalia J. Bayona-Vásquez, Travis C. Glenn, Jaime Gongora, Richard E. Green, Sally Isberg, Richard D. Stevens, David A. Ray

**Affiliations:** Department of Biological Sciences, Texas Tech University, Lubbock, TX, USA; Department of Environmental Health Science and Institute of Bioinformatics, University of Georgia, Athens, GA, USA; Sydney School of Veterinary Science, University of Sydney, Sydney, Australia; Department of Biomolecular Engineering, University of California, Santa Cruz, CA, USA; Centre for Crocodile Research, University of Sydney and Charles Darwin University, Sydney, Australia; Department of Natural Resources Management, Texas Tech University, Lubbock, TX, USA

**Keywords:** *Crocodylus porosus*, evolution, selection

## Abstract

Crocodilians are an economically, culturally, and biologically important group. To improve researchers’ ability to study genome structure, evolution, and gene regulation in the clade, we generated a high-quality *de novo* genome assembly of the saltwater crocodile, *Crocodylus porosus*, from Illumina short read data from genomic libraries and *in vitro* proximity-ligation libraries. The assembled genome is 2,123.5 Mb, with N50 scaffold size of 17.7 Mb and N90 scaffold size of 3.8 Mb. We then annotated this new assembly, increasing the number of annotated genes by 74%. In total, 96% of 23,242 annotated genes were associated with a functional protein domain. Furthermore, multiple non-coding functional regions and mappable genetic markers were identified. Upon analysis and overlapping the results of branch length estimation and site selection tests for detecting potential selection, we found 16 putative genes under positive selection in crocodilians, ten in *C. porosus* and six in *A. mississippiensis*. The annotated *C. porosus* genome will serve as an important platform for osmoregulatory, physiological and sex determination studies, as well as an important reference in investigating the phylogenetic relationships of crocodilians, birds, and other tetrapods.

## Introduction

Crocodilians (Order Crocodylia) are an ancient reptilian lineage whose extant members are likely to be among the most morphologically and genetically similar to the common ancestor of amniotes (Green, et al. 2014; Grigg 2015; Grigg, et al. 2001). Crocodilians and birds are the only extant members of the Archosauria, which also consists of the extinct lineages of dinosaurs and pterosaurs (Brusatte, et al. 2010). Within Crocodylia, the family Crocodylidae encompasses three genera of true crocodiles-*Crocodylus, Osteolaemus*, and *Mecistops*. They, along with the gharials (Gavialidae) are a sister clade to the third crocodilian family, Alligatoridae, the alligators and caimans (Brochu 2003; Densmore 1983). Crocodilians are important models for studies in phylogenetics (Brochu 2004, 1997; Gatesy, et al. 2003), osmoregulation (Grigg 2015), functional morphology (Rayfield and Milner 2008), sex determination (Deeming and Ferguson 1989; Lang and Andrews 1994; Pieau, et al. 1999; Western, et al. 1999), mating systems (Davis, et al. 2002; Lance, et al. 2009) and population genetics (Davis, et al. 2002; Ryberg, et al. 2002). Further, as they seem to possess an extremely effective immune system to combat pathogens that are abundant in their wild habitat (Jaratlerdsiri, et al. 2014; Merchant, et al. 2013; Merchant, et al. 2003), crocodilians are excellent models for understanding the evolution of the immune response. Knowledge of crocodilian genomes facilitates additional work in those areas and provides a key phylogenetic connection for studying relationships among amniotes and an opportunity to understand gene and genomic properties of extinct archosaurs.

In addition to the rationale presented above, recent analyses of whole crocodilian genomes suggest that they have evolved very slowly over the past several million years when compared to other tetrapods (Green, et al. 2014). Understanding the evolution, regulation and adaptive capabilities of the crocodilian genome and its genetic diversity can therefore provide information on how slow-evolving genomes manage to stay viable in the face of ever-changing environmental conditions.

Two annotated draft assemblies of the *Crocodylus porosus* genome are currently available. The first assembly, Cpor_2.0 (GCA_000768395.1; Green, et al. 2014), made use of Allpaths-LG (Gnerre, et al. 2011; Green, et al. 2014) to assemble data from Illumina short-insert and mate-pair libraries. The second assembly, CroPor_comp1, (GCF_001723895.1; Rice, et al. 2017) used Ragout (Kolmogorov, et al. 2014) to leverage a chromosome-scale alligator assembly with Cpor_2.0 to create a large-scale assembly for *C. porosus*. Although CroPor_comp1 has high contiguity, this is based on assumed orthologous contiguity. Thus, a high-quality, well-annotated *de novo* genome assembly of *C. porosus* similar in quality to the most recently released version of the *A. mississippensis* (Rice, et al. 2017) will allow a more comprehensive assessment of the species’ genome in terms of contiguity, gene space and annotations. This work attempts to bridge that gap by presenting an annotated and highly contiguous draft genome of *C. porosus*. We combined libraries available from the initial sequencing work (Green, et al. 2014) and included a *de novo in vitro* proximity-ligation Chicago library (Dovetail Genomics). Combining the Chicago library with Dovetail Genomics’ HiRise software pipeline, this assembly significantly reduces gaps in alignment originating from repetitive elements in the genome (Putnam, et al. 2016) and allows for increased confidence in gene predictions, thereby providing a vastly improved resource for researchers interested in crocodilian, archosaurian and vertebrate genomics.

In the new assembly, 23,242 genes were identified and annotated, improving markedly on the previous annotation. Repeat elements, microsatellite and tRNA annotations were also accounted for. All identified genes possessed an Annotation Edit Distance (*AED*) score of ≤ 0.3 in the MAKER2 pipeline (Holt and Yandell 2011), indicating high similarity with the provided transcript and protein evidence for *de novo* gene prediction and identification. Of the predicted genes, 96% were found to possess a functional protein domain as identified by InterProScan5 (ver.5.27-66) (Zdobnov and Apweiler 2001). Finally, using these data, a set of genes were identified that are likely under differential selection regimes, both in the alligator and crocodile lineages.

## Materials and Methods

### Library preparation and de novo shotgun assembly

The new improved assembly was generated using both raw reads from the previously released genome draft (Green, et al. 2014) and new Illumina sequencing data from a Chicago library prep from the same individual. Genomic DNA was isolated from a blood sample of a single male *C. porosus*, Errol, caught in the wild in the Northern Territory of Australia and currently housed at the Fort Worth Zoo (Texas, USA). Sequence data from three previous Illumina libraries with insert lengths of 167, 370, and 1800 bp (Green, et al. 2014) were trimmed and quality filtered using Trimmomatic (Bolger, et al. 2014), then assembled with Meraculous 2.0 (Chapman, et al. 2011) at Dovetail Genomics (Santa Cruz, CA, USA).

### Chicago library prep and scaffolding the draft genome

The Chicago library was prepared following methods from previous work (Putnam, et al. 2016) at Dovetail Genomics (Santa Cruz, CA, USA). Briefly, ≥ 0.5 μg of high molecular weight genomic DNA was used to reconstitute chromatin *in vitro* onto naked DNA and fixed with formaldehyde. Fixed chromatin was digested with *Dpn*II, resulting 5’ overhangs were filled in with biotinylated nucleotides, and free blunt ends were ligated together. After ligation, crosslinks were reversed and DNA was purified from proteins. Biotin that was not internal to ligated fragments was removed. DNA was sheared to a mean fragment size of ∼350 bp, and sequencing libraries were generated using NEBNext Ultra enzymes (New England Biolabs, Ipswich, MA, USA) and Illumina-compatible adapters. Biotin-containing fragments were isolated using streptavidin beads before PCR enrichment of the library. The Chicago library was sequenced on an Illumina HiSeq 2500 at HudsonAlpha to obtain PE150 reads.

Using Dovetail Genomics’s HiRise scaffolding pipeline, we mapped the shotgun data from the Chicago library to the draft input assembly obtained above, using a modified version of SNAP read mapper (http://snap.cs.berkeley.edu). We detected and omitted regions with abnormally high coverage for scoring joins and breaks. We analyzed the Chicago paired reads that mapped to the draft assembly to produce a likelihood model to identify putative misjoins and score prospective joins. Then, we filled gaps between contigs by scaffolding with the shotgun sequences from the Chicago library. We refer to this new *C. porosus de novo* genome as the Cpor_3.0 or Chicago-HiRise assembly (GenBank ID GCA_########).

### Comparison of C. porosus genome assemblies

There are currently three *C. porosus* assemblies, all generated using data from the same individual, Errol. For the first assembly, Illumina paired-end reads were generated from two short-insert libraries and one mate-pair library (Green, et al. 2014). The data were assembled with AllPaths-LG (Gnerre, et al. 2011). More recently, a highly-contiguous assembly from *Alligator mississippiensis* (Rice, et al. 2017) was used to re-scaffold the All-Paths assembly (Green, et al. 2014) using Ragout (Kolmogorov, et al. 2014). Here, we prepared and re-assembled using our Chicago libraries (see *Library preparation and de novo shotgun assembly*, above). Because all data originated from the same source, direct comparisons among assemblies can be made to detect the differences without the need to account for inter-individual variation.

We used the script ‘stats.sh’ from BBMap (sourceforge.net/projects/bbmap/) to calculate basic assembly statistics for all three assemblies. Next, we used BUSCO v 3.0.2 (Simao, et al. 2015) to obtain quantitative measures of gene content, using 3,950 single-copy orthologous genes from the tetrapod lineage database, tetrapoda_odb9, and setting chicken as the Augustus species gene finding parameter. We then used the JupiterPlot (https://github.com/JustinChu/JupiterPlot) pipeline to visually compare the assembly from (Rice, et al. 2017) (set as the reference) to our assembly, setting the minimum size of a contiguous region to render to 100 bp, considering all reference chromosomes larger than 100 bp, and using the largest reference scaffolds that are equal to 96.4% of our genome, to the full-length of the reference genome. Finally, MUMmer v. 4.0.0 (Kurtz, et al. 2004) was used to align and draw a dot plot to evaluate synteny between assemblies. For the synteny analysis using mummer we aligned 69 scaffolds from the reference that were larger than 1 Kb to 885 scaffolds from our assembly (query) that were larger than 1kb. The pairwise alignment of these scaffolds show some structural rearranges. For example, the longest scaffold in our query SciaK46_24 is 59,776,657 base pairs long and aligns to 54 different scaffolds from the reference. Similarly, the biggest scaffold from Rice et al. (2017) NW017728886.1, is 270,692,262 base pairs long and it aligns against 307 scaffolds from our query assembly. The average percent identity between scaffolds was 90.05%, with a minimum of 76.71% and a max of 100%. Additional quantitative elements from this comparative analysis can be found in Supplementary File S1.

### De novo gene annotation

Repeatmasker (Smit, et al. 1996) was run on the new genome assembly with *Crocodylus* as ‘–species’ option and the genome was soft masked. Gene annotation in the *C. porosus* was performed using MAKER2 pipeline (Holt and Yandell 2011), and SNAP (Korf 2004) was used as the *de novo* gene predictor. The MAKER2 pipeline was complemented with the transcript and protein FASTA files of *C. porosus* that were generated during the previous genome assembly annotation effort (Green, et al. 2014). This evidence was also used to train SNAP for more accurate gene prediction in the current assembly. General Feature Format (GFF) files with predicted gene models and FASTA files, one each for the transcript and corresponding protein sequences, were generated at the end of the MAKER2 run. Details of multiple options used in both runs of the MAKER2 pipeline, training of SNAP on the *C.porosus* genome, as well as details of re-running the MAKER2 pipeline with the trained SNAP program is described in the Supplementary Methods

### Post processing of annotations

Several steps were taken to generate the final functional annotation of the genes predicted from the MAKER2 run and were accomplished through multiple perl scripts provided with the MAKER2 package. Briefly, the ‘maker_map_ids’ was run to create a new ‘map file’ with revised nomenclature for the predicted genes in a numeric manner with a chosen prefix of ‘cPor’. Next, the scripts ‘map_fasta_ids’ and ‘map_gff_ids’ were executed on the FASTA and GFF files respectively along with the above ‘map file’ to update the previous nomenclature of predicted genes with the map file information. Finally, the ‘maker_functional_gff’ and ‘maker_functional_fasta’ were run to add putative BLAST (Altschul, et al. 1990) reports to the renamed GFF and FASTA files. In addition, the program InterProScan5 (v 5.27-66) (Zdobnov and Apweiler 2001) was used to add protein domains to the above annotated genes. The ‘ipr_update_gff’ script was used to add putative InterProScan5 results to the GFF and FASTA files.

### Identification of Microsatellites

To help anchor scaffolds from the Dovetail assembly with the previous linkage map for *C. porosus* (Miles, et al. 2009), the 282 *C. porosus* microsatellite loci present in GenBank (Supplementary File S2, Sheet 1) were screened using Repeatmasker v 4.0.5 and the masked file was aligned to the *C. porosus* genome using Burrows Wheeler Aligner (BWA) v 0.7.15 (Li and Durbin 2010). The resulting SAM file was converted into a binary file, sorted and indexed using Samtools v 1.3.1 (Li, et al. 2009) and the sorted alignment file was visualized against the *C. porosus* genome in Integrative Genomics Viewer (IGV) v 2.4.4. Thirty-four loci did not map to the genome, out of which 22 (KX055916.1 - KX055937.1) were allelic variants corresponding to a single locus Cj16. This locus was mapped to the genome with Cj16 primers (Isberg, et al. 2004) using the *in-silico* primer mapping algorithm in Geneious v 10.0.9 (Kearse, et al. 2012). The Cj16 forward and reverse primers mapped to a single region in contig SciaK46_869 and was included in the alignment file for further analysis. The alignment file was then analyzed to determine the relative distances between mapping positions.

Among the 248 remaining microsatellite loci that mapped to the genome assembly, 35 loci mapped to two or more positions within the same contig, 34 of which had a distance <900 bp and 27 of those 34 were <300 bp apart (Supplementary File S2, Sheet 3). On closer examination, we observed that these 34 loci had masked repeat sequences interspersed between two mapping positions. As a result, the first mapping position was selected for such loci and the other position was removed from further analysis. The remaining locus, (positions 26.7 Mbp apart) was excluded from the analysis. Seventeen GenBank IDs mapped to the same position in the genome as a previous locus and were considered duplicates of the first mapped locus and thus were removed from the analysis (Supplementary File S2, Sheet 4). Thus, in total, about 23% of the 282 microsatellite loci in GenBank were multi-mapping loci and were removed from further analysis. From the remaining loci, relative distances were calculated between the 131 adjacent loci mapping to the same contig and a distribution of these relative distances was constructed (Supplementary File S2, Sheet 5) in JMP^©^ Pro v.13 (SAS Institute Inc., Cary, NC).

### Identification of tRNAs

Transfer RNAs in the newly assembled *C. porosus* genome were predicted using tRNAscan-SE 2.0 (Lowe and Eddy 1997). The covariance model employed by tRNAscan-SE 2.0 was trained with training sets comprised of eukaryotic tRNAs. A subset of ten of the predicted tRNAs coding for amino acids were selected randomly and their sequences were searched against GtRNAdb (Chan and Lowe 2016) and tRNAdb (Juhling, et al. 2009). These sequences were found to be tRNAs predicted in a large number of other species in both the databases. Sequence and structure of the tRNAs were also provided by tRNAscan-SE 2.0 (Supplementary File S3, sheet 5 and sheet 6).

### Selection estimation by branch length

To identify genes potentially subjected to selection in one species or the other (*C. porosus* vs. *A. mississippiensis*), we conducted multiple tests of selection using orthologous genes. We considered including the gharial assembly, but it is of relatively poor quality compared to the *A. mississippiensis* and *C. porosus*, leading to multiple misalignments of orthologs and obviously incongruous branch length estimations. Consequently, we removed the gharial from our selection analyses. The current annotation of the chicken genome (GCA_000002315.3), was used as the outgroup for the analysis. ProteinOrtho v 5.16b (Lechner, et al. 2011) was used to identify single-copy orthologous genes in all three species.

Our first test was a per-gene branch length analysis. Orthologous amino acid sequences were aligned using MAFFT v 7.313 (Katoh, et al. 2005). TrimAl v 1.3 (Capella-Gutierrez, et al. 2009) was used to trim any unaligned ends, thereafter the alignments were converted to Phylip format using a custom python script. Then, PAML v 4.9g (Yang 2007, 1997) was used to calculate branch lengths for each alignment of orthologous genes from *C. porosus* and *A. mississippiensis* using the species tree ‘(alligator,crocodile),outgroup’ for each gene. Multiple custom bash and python scripts were used to parse input/output files when implementing the above steps (Supplementary File S4). For PAML specifically, we used the AAML package of PAML as we used the amino acid codon sequences for alignment and analysis purposes here. Once branch lengths for the amino acid sequences were estimated using PAML, we sorted branch lengths (based on branch length values) of *C. porosus* and *A. mississippiensis* using chicken as outgroup. The log-transformed ratios of the *C. porosus* to the *A. mississippiensis* branch lengths for each gene were used to infer genes under potential selection in *C. porosus*, while the ratios at the other end of the range implied genes under potential selection in *A. mississippiensis*. The top 2.5% of the genes, for each species were considered for further analysis of their functional significance.

### Statistical tests for adaptive evolution of codons (site-specific model)

An additional statistical test for adaptive evolution using the site-specific model of CODEML (from the PAML v4.9g package, (Yang 2007)) was also performed. This allowed us to identify potential genes under positive selection by analyzing the dn/ds (non-synonymous substitution to synonymous substitution) ratios of the genes in *C. porosus* using chicken as outgroup. Briefly, the program pal2nal (Suyama, et al. 2006) was used to analyze the species specific protein-coding sequences (CDS) and the aligned protein sequences (generated previously through MAFFT and trimAl) to create aligned CDS sequences for the two crocodilian species as well as for the chicken outgroup. A custom perl script and BEDtools (Quinlan and Hall 2010) was used to extract all CDS sequence from GFF files of *C. porosus* (Supplementary File S4). Multiple custom python and bash scripts were utilized to generate files in an acceptable format for CODEML (Supplementary File S4). The models selected in CODEML were M0, M1, M2, M7 and M8 for site selection to test adaptive evolution of genes (Anisimova, et al. 2001; Swanson, et al. 2001; Yang and Bielawski 2000; Yang and Nielsen 2002). M0 implies the null model while M1, M2, M7 and M8 are alternative models that were used in a likelihood ratio test to identify sites-specific selection in the species. Statistical significance of the difference of log-likelihood values over the χ^2^ distribution table was used to identify genes potentially evolving under positive selection. Details of all programs and options used in this gene selection analysis (branch length ratio comparison and site selection models) can be found in Supplementary Methods.

### Detecting codon evolution using the branch-site model

For branch-site model tests, we used the same crocodile, alligator, and chicken datasets from the aforementioned analyses. PAML’s branch-site model test has demonstrated robustness when analyzing species with extreme divergences (Gharib and Robinson-Rechavi 2013). We therefore incorporated an additional four species into our analyses: pigeon (Rock Pigeon - *Columba livia* - GCA_001887795.1), barn swallow (Barn swallow - *Hirundo rustica* - GCA_003692655.1), brown kiwi (Brown Kiwi - *Apteryx australis* - GCA_001039765.2), and common box turtle (*Terrapene carolina* - GCA_002925995.2). Single copy orthologous gene regions were curated from all seven species using ProteinOrtho v 5.16b (Lechner, et al. 2011) and trimmed using TrimAl v 1.3 (Capella-Gutierrez, et al. 2009). Amino acid sequences were then converted to codon alignments using pal2nal (Suyama, et al. 2006). Each shared single-copy orthologous gene alignment was used to construct a maximum likelihood tree using RAxML v 8.2.11 with 1000 bootstrap iterations(Stamatakis 2014). An unrooted species tree was created from the resulting single-copy orthologous gene trees using ASTRAL-III v 5.6.3 (Zhang, et al. 2018).

To detect positively selected genes, two separate datasets were generated with the alligator and crocodile each serving in the foreground position on the phylogeny. We applied PAML’s branch-site model to detect signatures of selection along specific branches with model M2a (selection) and NSsites = 2. We compared the null model (codons evolve under purifying or neutral selection, fix ω = 1 & ω = 1) against the alternative model (codons under positive selection, fix ω = 0). Likelihood ratio statistics were calculated for each branch-site model of an orthologous gene by CODEML. Significance (p < 0.05, df = 1) of the log likelihood ratio statistic comparisons was evaluated against a χ^2^ distribution. Additionally, a Bonferroni correction was applied to the log likelihood ratio statistics.

### GO-Enrichment of genes under positive selection and identification of gene network pathways

Once the single-copy orthologous genes under putative positive selection were identified by the methods above, we identified overlaps. Although there were no genes that overlapped all three selection approaches, 16 genes were identified by both the from the site-selection and branch-estimation methods. We analyzed these 16 genes for GO term enrichment to understand if they were involved in particular cellular and metabolic pathways. The amino acid FASTA sequences for all 16 genes were used as input in the KOBAS 3.0 program (KEGG Orthology Based Annotation system) (Wu, et al. 2006; Xie, et al. 2011).The result generated the list of input genes enriched for their associated GO-terms by employing the hypergeometric test / Fisher’s exact test for statistical analysis and the Benjamini and Hochberg method of multiple test correction (Benjamini and Hochberg 1995).

## Results and Discussion

Our *de novo* assembly represents a significant improvement compared to the initial *de novo* assembly using Allpaths-LG (Green, et al. 2014) (Table 1). While the total length of the assembly remained similar for both *de novo* methods (Allpaths-LG and Chicago-HiRise), statistics improved by 86-fold for scaffold N50 and 74-fold for scaffold N90 when using information from the Chicago libraries. Consequently, the total number of scaffolds was reduced by ∼90%. Such improved contiguity is expected to increase our ability to identify genes in the assembly and this was indeed the case (Table 2). Although the current assembly had lower contiguity than the Ragout reference-based assembly of (Rice, et al. 2017), our Chicago-HiRise assembly is based entirely upon *de novo* analyses and does a superior job in assembling genes (Table 2).

**Table 1.**
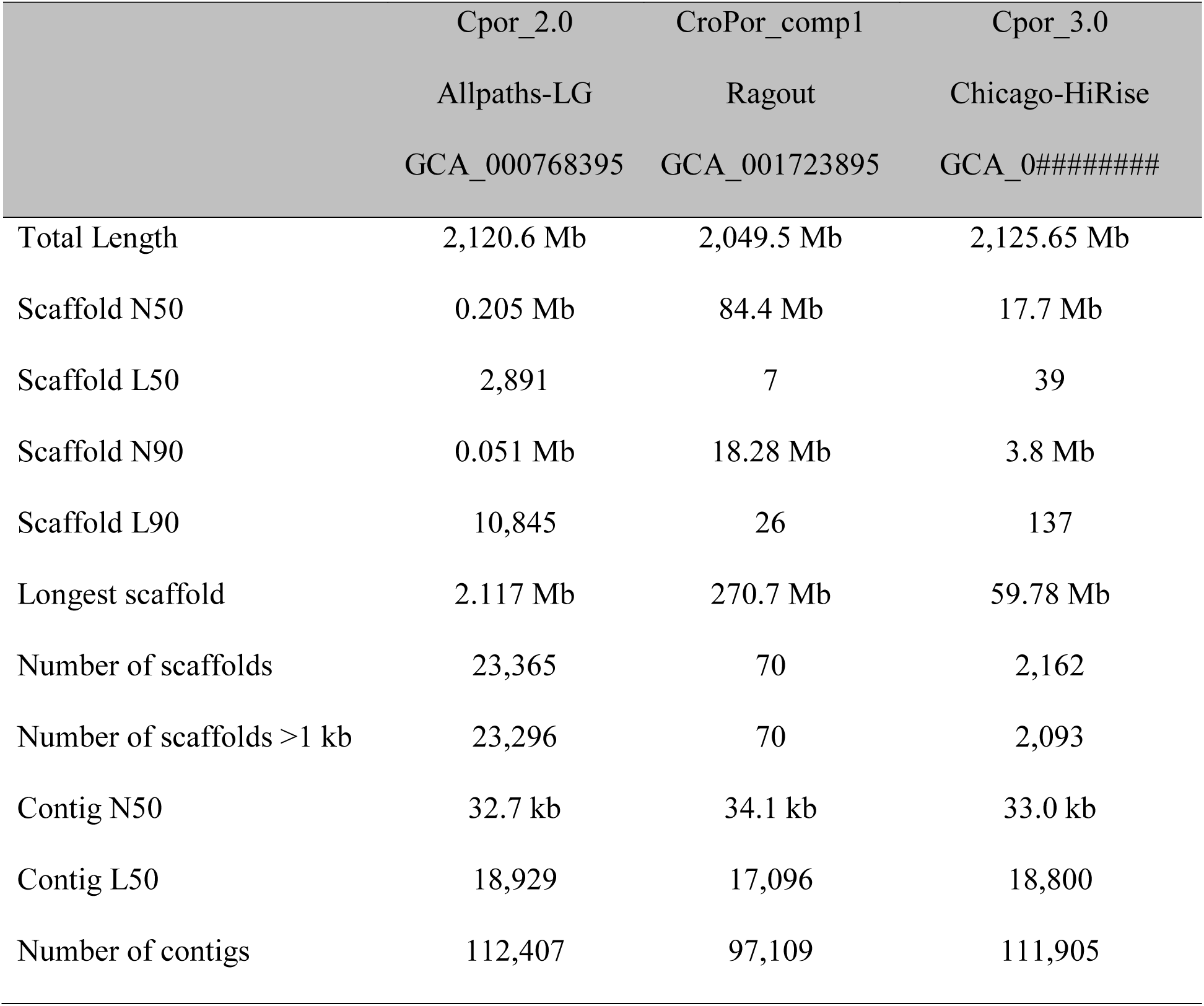
Quality statistics for available assemblies of *C. porosus*, including our draft and the current HiRise assembly

**Table 2.** BUSCO summary stats when searching for 3,950 orthologous genes from tetrapods. The percentage of genes relative to the total in the database are given in parentheses.

|  | **Cpor_2.0** | **CroPor_comp1** | **Cpor_3.0** |
| --- | --- | --- | --- |
| Total BUSCOs (comp+frag) | 3723 (94.3) | 3682 (93.2) | 3808 (96.4) |
| Complete single copy | 3338 (84.5) | 3435 (87.0) | 3535 (89.5) |
| Complete duplicated | 22 (0.6) | 20 (0.5) | 23 (0.6) |
| Total Complete BUSCOs | 3360 (85.1) | 3455 (87.5) | 3558 (90.1) |
| Fragmented BUSCOs | 385 (9.7) | 247 (6.3) | 250 (6.3) |
| Missing BUSCOs | 205 (5.2) | 248 (6.2) | 142 (3.6) |

### Comparison of C. porosus genome assemblies

The *de novo* assemblies were similar in overall size and GC content, but the contiguity of the Chicago-HiRise assembly was much better (Table 1). The Ragout assembly differed slightly in base composition and contained more than twice as many N’s (>5% vs. <2%). In general, our Chicago-HiRise assembly presents intermediate values of contiguity between the Ragout assembly of (Rice, et al. 2017) and the Allpaths-LG assembly of (Green, et al. 2014). Our Chicago-HiRise assembly is the longest among all (2,125.65 Mb), representing an increase of 5.08 Mb over the AllPaths-LG assembly and of 76.11 Mb over the Ragout assembly. An analysis of the raw reads from Green et al. 2014 using Kmergenie v.1.7044 (Chikhi and Medvedev 2014) yielded an estimated size of 2,089.69 Mb, a value in good agreement with our assembly size.

Our Chicago-HiRise assembly yielded the highest count of total BUSCOs (Table 2), from single-copy, duplicated genes, and fragmented genes, when compared to the other assemblies. It also had the lowest number of missing BUSCOs (142, 3.6%) among all. This indicates that our assembly has the best representation of gene space for *C. porous*. Increasing the length by 0.24% and 3.7% allowed an increase of 2.1% and 3.2% of BUSCO matches, when compared to the Allpaths-LG and Ragout assemblies, respectively. This pattern is shown in the Jupiter Plot, where mostly small scaffolds in our Chicago-HiRise assembly are not represented in the Ragout assembly (Fig. 1A) and very few and small translocations are detected. Similarly, in the MUMmer alignment and dot plot, we found high synteny with very minor syntenic discontinuities (Fig 1B).

**FIGURE 1.**
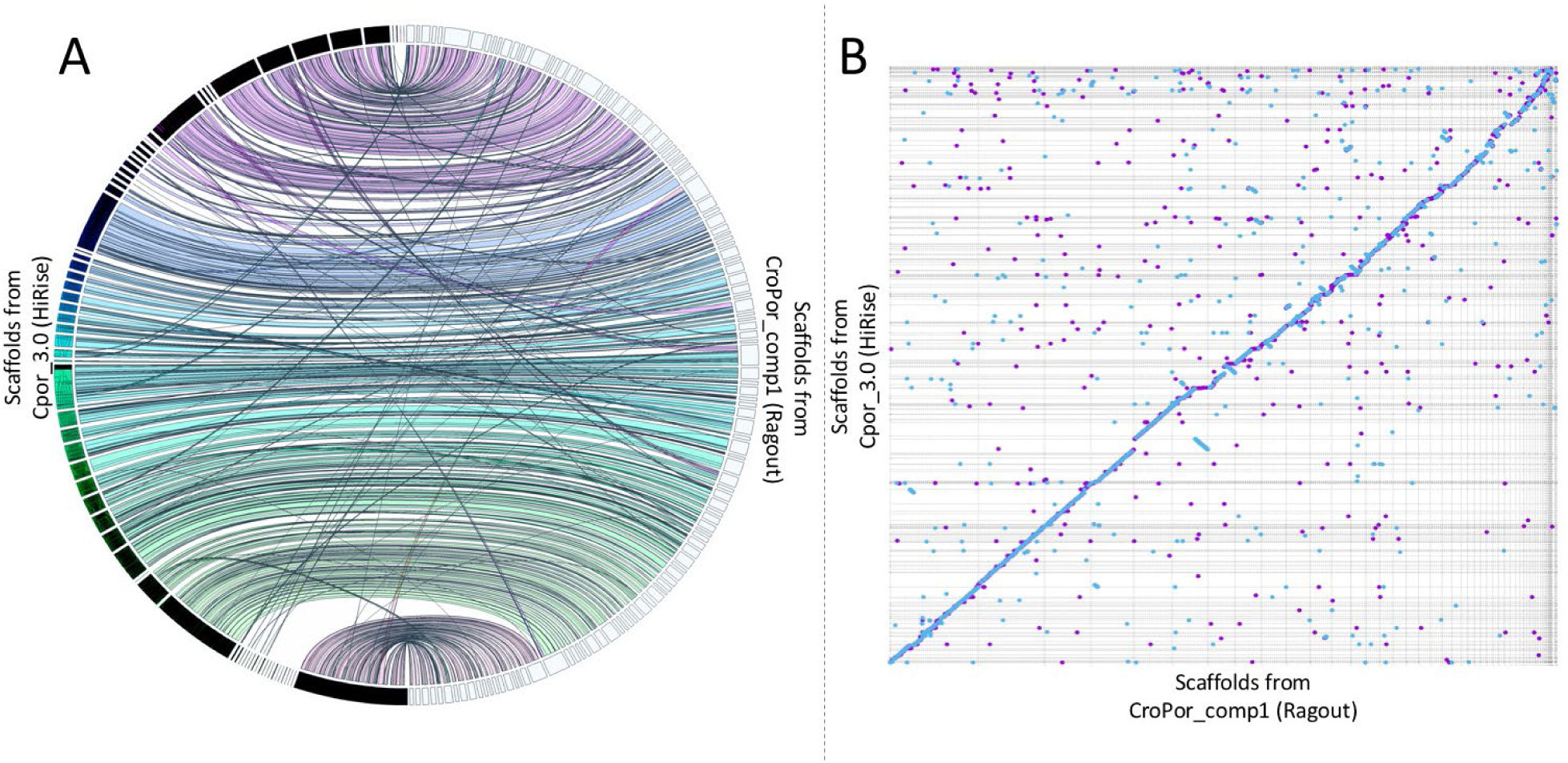
Synteny analyses between our Chicago HiRise assembly and the highly contiguous Ragout assembly from Rice et al. (2017). A) Jupiter Plot of correspondence between assemblies considering the total length of both reference and query genomes. B) Dot plot (MUMmer plot) of the percent identity in the alignment generated by MUMmer. The blue dots along the slope demonstrates that both assemblies are highly colinear. Blue dots represent forward matches and purple dots represent reverse matches.

### De novo gene annotation

A total of 23,242 genes were predicted and annotated in our Chicago-HiRise assembly compared to 13,321 genes in the initial AllPaths-LG assembly. From the 23,242 annotated genes, 22,226 genes (96%) were associated with one or more functional domains as identified by InterProScan5. One example representing a single gene along with its identified sections as predicted by MAKER2 pipeline with the integrated and trained SNAP gene predictor is shown in Supplementary File S5. A total of 7,155 unique genes were identified with GO annotations (Supplementary File S6).

Each annotated gene was assigned an *AED* score ranging from 0 to 0.3, where 0 indicated a perfect match between the intron and exon coordinates of an annotation and its aligned evidence. A distribution of the number of genes with their corresponding *AED* scores as identified in this study in *C. porosus* (Fig 2) and illustrates close resemblance of the genes with the provided transcript and protein evidence.

**FIGURE 2.**
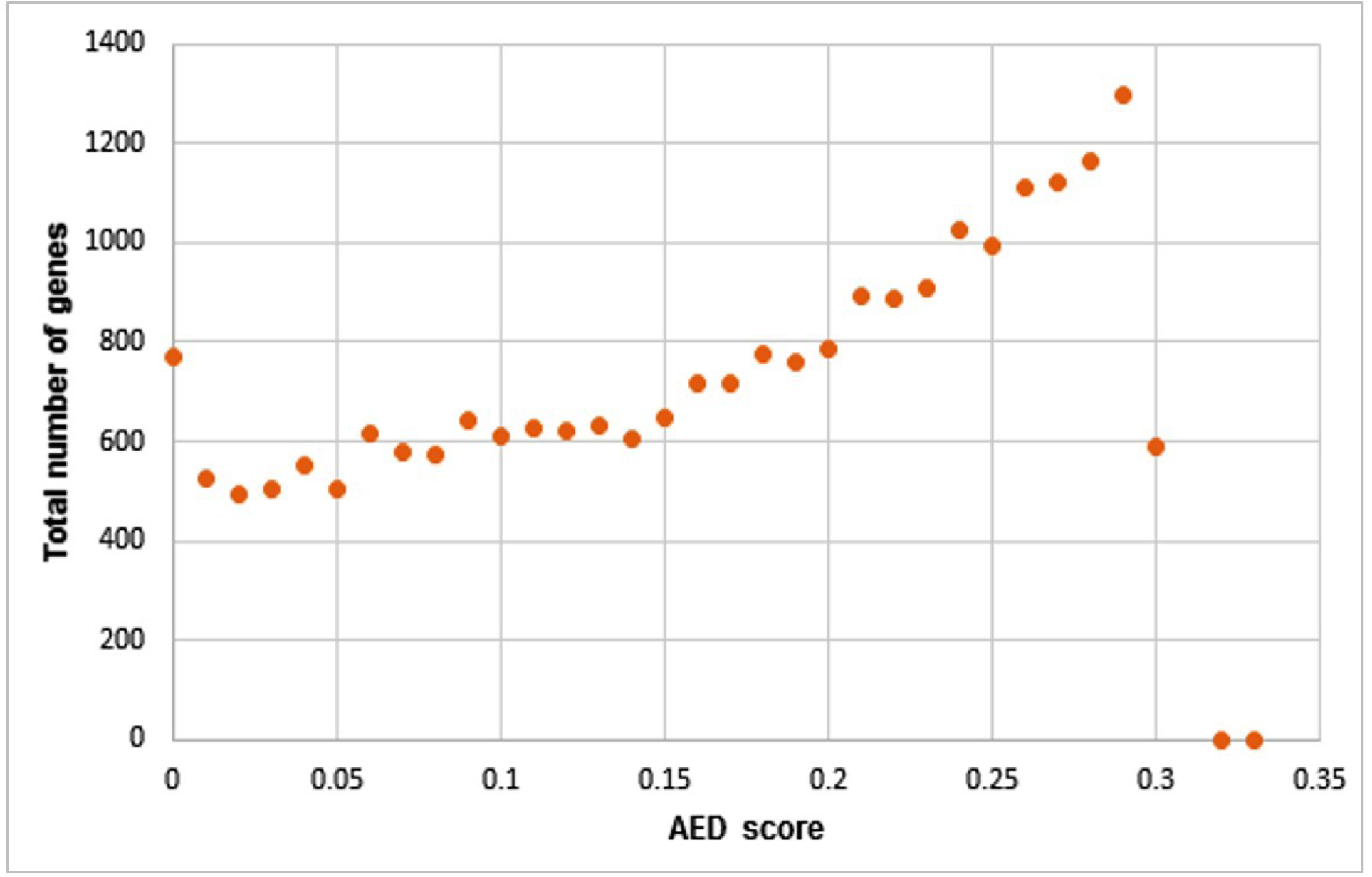
Representation of total number of unique genes as percentage of their corresponding Annotation Edit Distance (AED) scores as analyzed by MAKER2 pipeline form the *C. porosus* genome assembly.

### Microsatellite identification

The alignment of microsatellites of *C. porosus* to the genome assembly confirmed the loci to be scattered throughout the genome and unlikely to be subject to linkage. Of the 282 microsatellite loci aligned, 34 did not map to the genome, 155 mapped uniquely to a single location and 93 mapped to two or more loci. The relative distances between 131 adjacent loci mapping to the same contig are presented in Supplementary File S2, Sheet5. Twelve of these distances are < 960 kb, and all others are > 1 Mb apart. Ten of the distances were greater than 10 Mb apart. The locations of these microsatellite loci can be used in future studies to verify linkage via pedigree analyses (Miles, et al. 2009), and to order and orient scaffolds along chromosomes.

### tRNA prediction and identification

A total of 1,211 tRNAs were detected by tRNAscan-SE 2.0, out of which 437 were tagged as pseudogenes characterized by the absence of confirmed primary or secondary structures. These pseudogenes usually have low Infernal as well as Isotype bit scores in the predicted output (Supplementary File S3). In total, 16 tRNAs were found to have undetermined isotypes and 134 tRNAs were chimeric, with mismatched isotypes. There were 619 tRNAs coding for the twenty standard amino acids, and five tRNAs were found to code for selenocysteine. Among all tRNAs identified, 93 tRNAs harbored introns, out of which 32 were predicted to be pseudogenes, seven were chimeras. No suppressor tRNAs were identified in the analysis.

### Branch length analysis of selection

Given that crocodilians have relatively slow overall genome evolution (Green, et al. 2014), and because *A. mississippiensis* and *C. porosus* have different habitat preferences, we sought to identify genes potentially evolving under strong positive selection. We compared branch lengths of gene pairs in *C. porosus* and *A. mississippiensis*. A histogram shows the distribution of the log-transformed branch length ratios in these two crocodilians (Fig. 3). All genes potentially under differential selection in *A. mississippiensis* and *C. porosus* under the branch length estimation analysis are given in Supplementary Files S7-S8. Under a model of neutral evolution, few genes were identified as evolving under positive selection. We identified 47 genes potentially evolving under positive selection in *C. porosus* and 41 in *A. mississippiensis* respectively. These genes represent candidates for differential selection and rapid evolution in one crocodilian but not the other.

**FIGURE 3.**
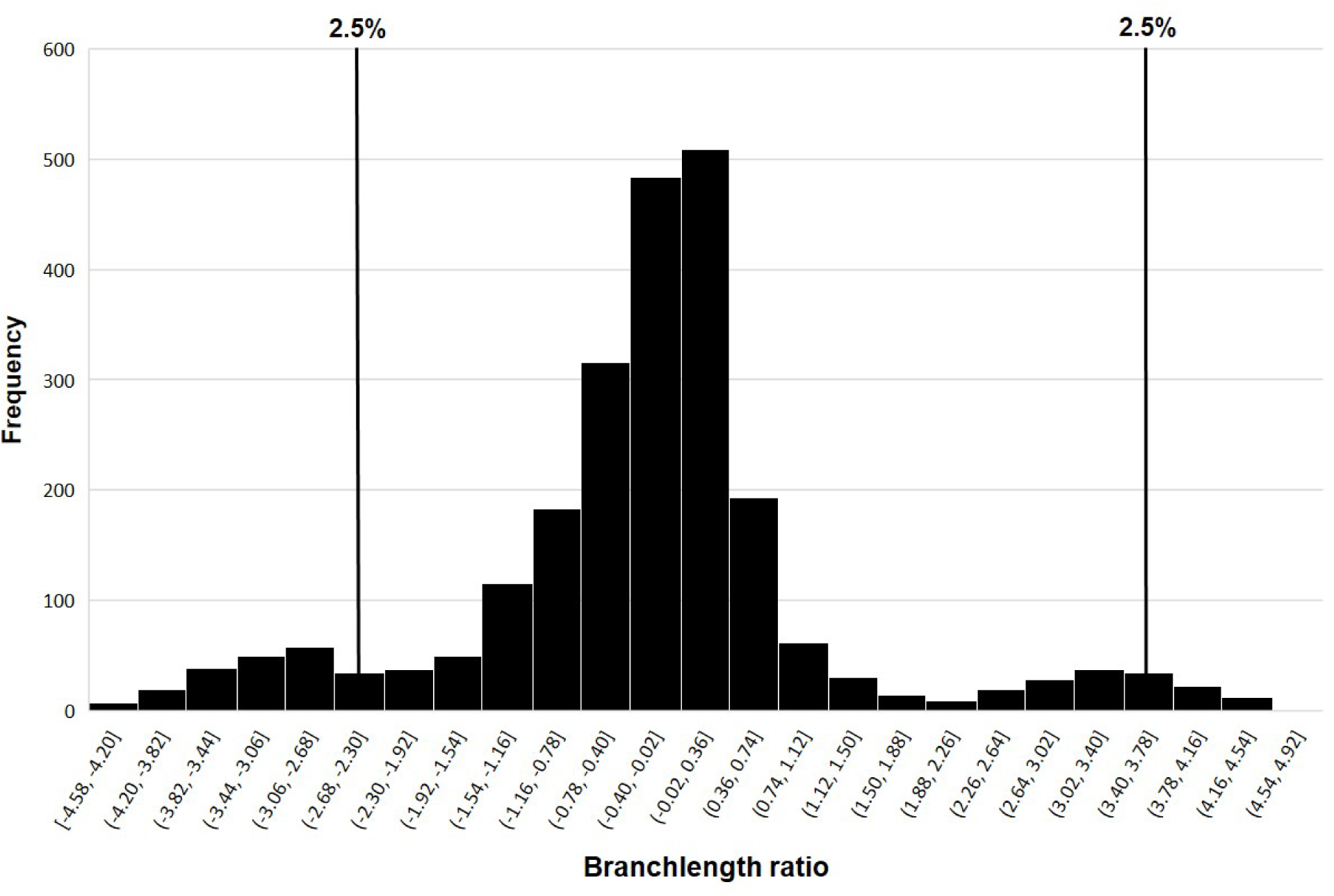
Histogram of the branch length ratio of *A. mississippiensis* and *C. porosus* with chicken as the outgroup. The two tails of the histogram correspond to the 2.5% of the genes in the *A. mississippiensis* and *C. porosus* respectively that are under potential selection. Vertical lines indicate the 2.5% cutoff limits in the histogram.

Genes identified as subject to rapid change under the branch length analysis in *C. porosus* included those directly or indirectly involved in salt metabolism and sodium transport. Some of these genes included the Na^+^-Ca^++^ exchanger/integrin-beta4 protein, sodium/calcium exchanger protein, peroxiredoxins and dehydrogenases membrane proteins – that are related to peroxide and free radical scavenging, increasing due to increased ionic and osmotic stress (salt stress) and can degrade hydrogen peroxide to water. As would be expected based on differences in natural history between *C. porosus* and *A. mississippiensis*, the above genes were absent from the list of rapidly evolving genes in *A. mississippiensis*. For example, given the alligator’s comparatively strict restriction to freshwater habitats one would not expect to find positive selection in osmoregulatory genes. Instead, heat shock genes (HSP40/DnaJ peptide-binding) were prominent in the *A. mississippiensis* list. Heat shock proteins are known to be upregulated in cold stress (Colinet, et al. 2010; Rinehart, et al. 2007; Stetina, et al. 2015) and alligators are known to have much higher cold tolerance as compared to crocodiles. A list of the genes with their putative functions in both *C. porosus* and *A. mississippiensis* can be found in Supplementary File S7-S8, respectively.

### Site model tests for selection

A total of 2357 single-copy orthologous genes were identified for *C. porosus, A. mississippiensis*, and the chicken (outgroup). The dn/ds ratio is an effective measure of the strength of natural selection acting on protein coding genes (dos Reis and Wernisch 2009; Sharp, et al. 2005). This ratio is indicative of which genes are evolving neutrally (dn/ds =1); are under negative or purifying selection (dn/ds < 1) as well as for ones that are being acted on in an adaptive or diversifying manner (positive selection; dn/ds > 1). The majority of the protein coding genes will have conserved codons and will probably not undergo positive selection (dos Reis and Wernisch 2009; Sharp, et al. 2005). This is because majority of protein-coding sequences are better adapted for functionality and changes will not necessarily lead to selective advantage (Hughes 1999). Of the 2,357 orthologous genes using the chicken outgroup, the vast majority (93.5%) exhibited a value of dn/ds < 1, while 387 orthologous genes (∼ 16%) exhibited signs of positive selection (Fig. 4). Genes involved in membrane pore channel transport, sodium bicarbonate cotransporter, sodium/hydrogen exchanger, sodium-phosphate symporter, sodium/potassium/calcium exchanger, Amiloride-sensitive sodium channel subunit gamma, sodium/potassium gated channel protein, SLC-mediated transmembrane transport, heat shock proteins, DNA repair, chondroitin sulfate biosynthesis were some of the many that were identified. Details of this list of 387 genes identified through the site selection procedure can be found in Supplementary File S9.

**FIGURE 4.**
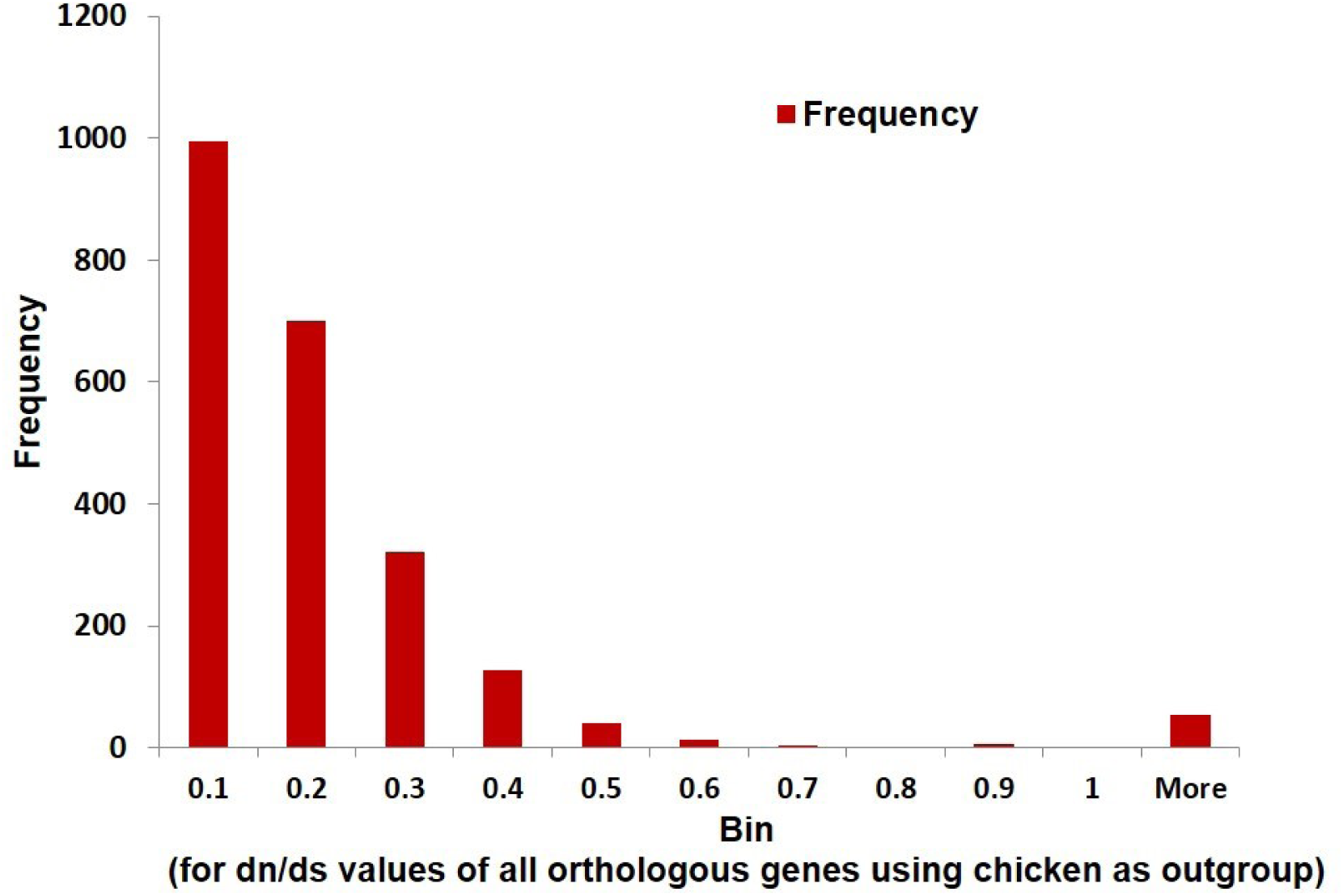
Histogram of dn/ds values for all genes of *C. porosus* using the M0 model with *A. mississippiensis*. A majority of the 2,357 single-copy orthologous genes are expectedly under purifying selection.

### Branch-site model testing

The branch-site model analysis yielded 17 genes under positive selection (Supplementary Table 1). Interestingly, the protein ATP1A1 was identified with high statistical support (p < 0.0005) for crocodile. ATP1A1 encodes an alpha-1 subunit for the cation transporter ATPase which is responsible for maintaining the electrochemical gradients across plasma membranes. As primarily marine inhabitants, crocodiles depend on effective osmoregulation to maintain ionic homeostasis. Crocodiles are unique compared to alligators as they have lingual salt glands to assist in excreting excess sodium and chloride (Cramp, et al. 2008). The necessity for efficient salt excretion could explain why ATP1A1 gene exhibited a strong signal of positive selection under the branch-site model of PAML.

We overlapped the findings of the three selection approaches applied in this study. While the overlap of all three approaches did not find commonalities, the overlap of the branch length estimation approach and the site selection approach resulted in 16 single-copy orthologous genes. Ten of the 16 belonged to *C. porosus*, and six to *A. mississippiensis*. The identification of these loci in *C. porosus* and *A. mississippiensis* could be starting points to investigate the biological differences associated with the salt-tolerance evolution in each species as well as habitat preferences. For example, *C. porosus* is known to be seagoing within tropical climates, whereas *A. mississippiensis* rarely leaves freshwater and has a range that includes temperate to semi-tropical climates. The details of these 16 genes are represented in Table 3.

**Table 3.** List of 16 genes under potential selection (and overlap of two selection tests) in *C. porosus* and *A. mississippiensis*

| Query | Species | Abbreviated | Annotation |
| --- | --- | --- | --- |
| amisp005461 | *A. mississippiensis* | CHST7 | Carbohydrate sulfotransferase 7 |
| amisp005516 | *A. mississippiensis* | HOXC13 | Homeobox protein Hox-C13a-like |
| amisp016775 | *A. mississippiensis* | IFT122 | Intraflagellar transport protein 122 homolog isoform X1 |
| amisp017613 | *A. mississippiensis* | JMJD4-1 | jmjC domain-containing protein 4 |
| amisp032461 | *A. mississippiensis* | RNF126 | E3 ubiquitin-protein ligase RNF126 |
| amisp034033 | *A. mississippiensis* | POLDIP3 | Polymerase delta-interacting protein 3 |
| cPor_01965-RA | *C. porosus* |  | Zinc finger protein 143 |
| cPor_06982-RA | *C. porosus* | GALK1 | Galactokinase |
| cPor_09447-RA | *C. porosus* | CTBP2 | C-terminal-binding protein 2-like isoform X1; Belongs to the D-isomer specific 2-hydroxyacid dehydrogenase family |
| cPor_11586-RA | *C. porosus* | KCNK10 | Potassium channel subfamily K member 10; Belongs to the two pore domain potassium channel (TC 1.A.1.8) family |
| cPor_15403-RA | *C. porosus* | TLX1-1 | T-cell leukemia homeobox protein 1 |
| cPor_15737-RA | *C. porosus* | SLC25A17 | Peroxisomal membrane protein PMP34; Belongs to the mitochondrial carrier (TC 2.A.29) family |
| cPor_15755-RA | *C. porosus* | NPTXR | Neuronal pentraxin receptor |
| cPor_18091-RA | *C. porosus* | GRAMD2 | GRAM domain-containing protein 2 |
| cPor_18867-RA | *C. porosus* | HNRNPLL | Heterogeneous nuclear ribonucleoprotein L-like |
| cPor_19471-RA | *C. porosus* | UBXN2B | UBX domain-containing protein 2B |

### GO-term enrichment for genes and potential gene networking pathways in *C. porosus*

For the overlapping 16 genes identified to be under positive selection from both the branch length estimation and site selection models, we identified 61 different GO enrichment categories. However, only 12 of these 61 categories had p-corrected values <0.05. As expected from independent analysis of the two selection methods, some of the common categories among these 12 included genes involved in chondroitin sulfate biosynthesis pathway (later stages), RNA polymerase III transcription initiation, carbohydrate metabolism as well as pore domain potassium channels. In addition, 205 GO terms (also selected based on p-corrected < 0.05) were identified in the same analysis. Some of the prominent GO terms in which these 16 genes were enriched included limb development, metabolism, chondroitin sulfotransferase activity, multiple membrane transporter activity proteins, nail development, tongue morphogenesis and potassium ion leak channel activity. All the above function/categories are of functional significance in members of Crocodylia thus reinforcing the rationale of our gene enrichment analysis. The details of the analyses can be found in Supplementary File S10.

To analyze a putative gene networking present in these potentially evolving genes in the *C. porosus*, and *A. mississippiensis*, we analyzed the amino acid sequences on the REACTOME server v.69 (Croft et al. 2013). While REACTOME typically maps the query inputs to their highly curated human database to analyze gene networks/pathways, we used the option of “species comparison” (with input as chicken) when performing the analysis. This helped analyze the input (crocodilian) query against the human database that are only orthologous in sequences to the chicken. The gene networking pathway (Fig.5) revealed transport of small molecules, vesicle-mediated transport, signal transduction, metabolism, DNA replication and few others. Expectedly, these matched with the nature of the 16 genes in context as well as with their associated GO terms. Thus, the annotated genes, their associated GO terms and corresponding enrichment analysis on KOBAS 3.0 and finally the gene networking information from REACTOME helped us establish a comprehensive idea of the type of crocodilian genes under potential selection and evolving rapidly in both the *C. porosus* and *A. mississippiensis.*

**FIGURE 5.**
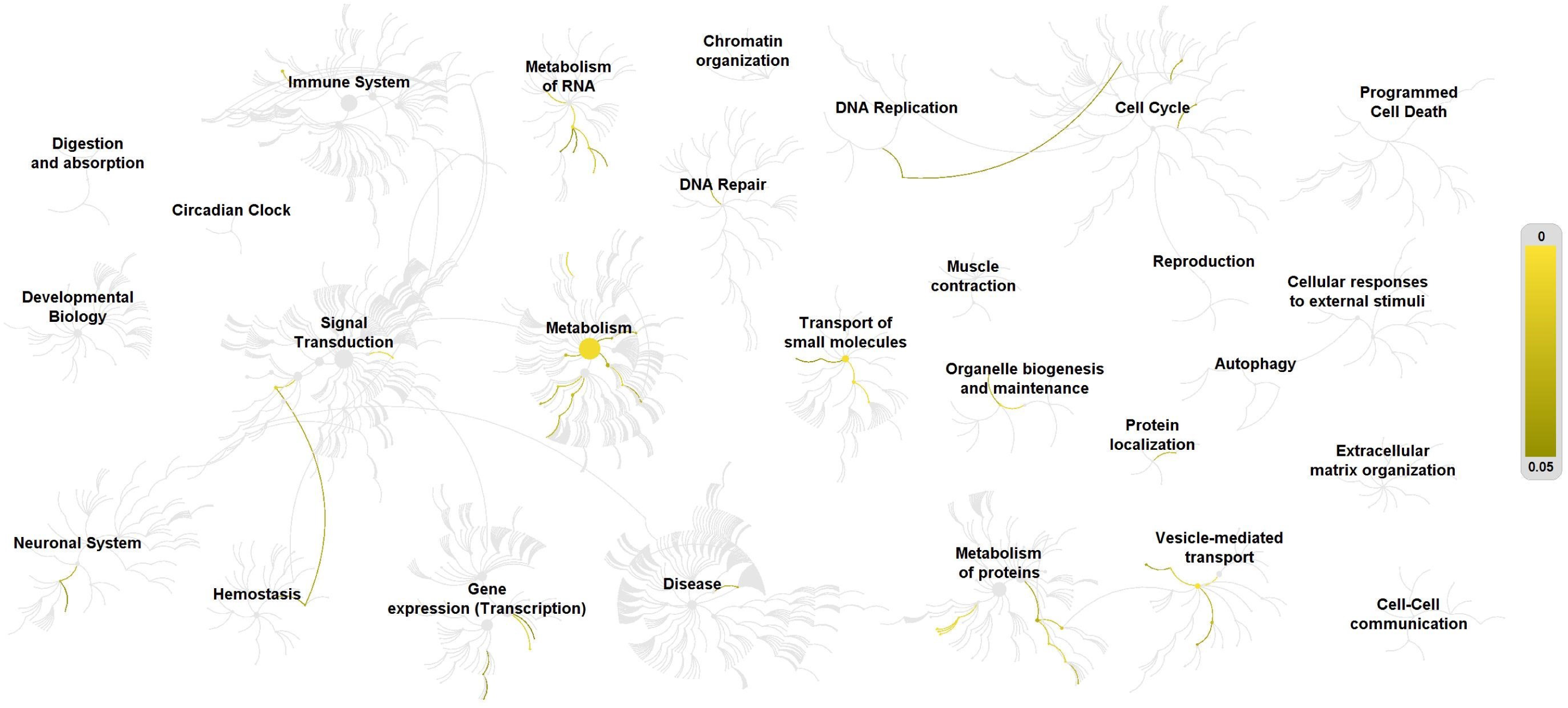
Representation of gene networking pathways for 16 genes found in *C. porosus* and *A. mississippiensis* that are under potential selection. Analysis was performed in REACTOME (v. 69) with *G. gallus* and *H. sapiens* as ortholog species comparison. The networking pathways signify interacting genes and pathways as predicted from the 16 input genes. The yellow color gradient (intensity) corresponds to a probability of overlap of the query genes with that of the gene networking pathways on the REACTOME server. Darker colors signify a higher probability of overlap (closer to p = 0.05), while a lighter yellow signifies a lower probability of overlap with a networking pathway (p = 0).

## Conclusion

A highly contiguous genome *de novo* assembly was constructed based on Illumina short read data from paired end and *in vitro* proximity-ligation Chicago library. The new assembly exhibits improved scaffold lengths over the AllPaths-LG assembly (Green, et al. 2014) and better assembly of genes and assessments of genome space occupancy when compared to the (Rice, et al. 2017) assembly. We identified 23,242 genes with 96% of those genes possessing a functional domain and 7,155 unique genes were associated with one or more GO terms, also an improvement relative to the AllPaths-LG and Ragout assemblies. We identified 1,211 tRNA and 155 previously characterized microsatellites mapped uniquely to a single location in the genome while 93 microsatellites mapped to multiple genomic locations. Multiple selection tests showed genes in both *C. porosus* and *A. mississippiensis* under potential positive selection. The enrichment of genes in certain cellular and metabolic pathways such as potassium channel pore domain protein and peroxisomal membrane proteins make sense due to the natural habitat of *C. porosus* and their adaptations to the saline environment. Additionally, the rapid directional evolution of heat shock proteins in *A. mississippiensis* is consistent when considering the higher cold tolerance of alligators relative to crocodiles and all other crocodilians. It might be noted here that the potentially high number of orthologous genes under positive selection when analyzed through the site model (387) could be partially due to the very low number of species used for the analysis.

With no other highly-contiguous crocodilian genome-assembly at our disposal, we could only use the *A. mississippiensis* assembly (the gharial assembly being of very low quality, was left out) along with two outgroups for the phylogenetic analyses. It is our hope that as more well-annotated genomes of other crocodilians are generated, subsequent phylogenetic analyses will be more comprehensive. Finally, with a highly contiguous and well-annotated genome assembly of *C. porosus*, a number of fields may benefit. The genome may serve as a resource for mapping comparative phylogenetic traits in sister crocodilians as well as defining novel phylogenetic relationships of birds. The newly annotated *C. porosus* genome assembly, Cpor_3.0, can also provide a robust platform for investigations in osmoregulatory research, functional morphology, as well as sex determination studies.

## Acknowledgements

This work was supported by the National Science Foundation (MCB-0841821, DEB-1020865, and DEB-1838283 to D.A.R.) and Rural Industries Research and Development Corporation (PRJ-005355 and PRJ-002461 to S.I. and J.G.). A University of Sydney Bridging Support grant to Jaime Gongora also supported the project. We thank the staff of Dovetail Genomics for help in preparing and processing the Chicago library and HiRise assemblies. The High Performance Computing Center (HPCC) at Texas Tech University and the Georgia Advanced Computing Resource Center at the University of Georgia provided computational infrastructure and technical support throughout the work. Additional support was provided by the College of Arts and Sciences at Texas Tech University.

## Notes

http://myweb.ttu.edu/daray/Pubs_files/Supplement_v24_GBE.zip

